# Long axial-range double-helix point spread functions for 3D volumetric super-resolution imaging

**DOI:** 10.1101/2024.07.31.605907

**Authors:** Yuya Nakatani, Scott Gaumer, Yoav Shechtman, Anna-Karin Gustavsson

**Author notes:** **Corresponding Author** Anna-Karin Gustavsson – Department of Chemistry, Department of Biosciences, Department of Electrical and Computer Engineering, Smalley-Curl Institute, and Center for Nanoscale Imaging Sciences, Rice University, Houston, Texas 77005, United States; Department of Cancer Biology, The University of Texas MD Anderson Cancer Center, 1515 Holcombe Blvd, Houston, Texas 77030, United States.

## Abstract

Single-molecule localization microscopy (SMLM) is a powerful tool for observing structures beyond the diffraction limit of light. Combining SMLM with engineered point spread functions (PSFs) enables 3D imaging over an extended axial range, as has been demonstrated for super-resolution imaging of various cellular structures. However, super-resolving structures in 3D in thick samples, such as whole mammalian cells, remains challenging as it typically requires acquisition and post-processing stitching of multiple slices to cover the entire sample volume or more complex analysis of the data. Here, we demonstrate how the imaging and analysis workflows can be simplified by 3D single-molecule super-resolution imaging with long axial-range double-helix (DH)-PSFs. First, we experimentally benchmark the localization precisions of short- and long axial-range DH-PSFs at different signal-to-background ratios by imaging of fluorescent beads. The performance of the DH-PSFs in terms of achievable resolution and imaging speed was then quantified for 3D single-molecule super-resolution imaging of mammalian cells by DNA-PAINT imaging of the nuclear lamina protein lamin B1 in U-2 OS cells. Furthermore, we demonstrate how the use of a deep learning-based algorithm allows the localization of dense emitters, drastically improving the achievable imaging speed and resolution. Our data demonstrate that using long axial-range DH-PSFs offers stitching-free, 3D super-resolution imaging of whole mammalian cells, simplifying the experimental and analysis procedures for obtaining volumetric nanoscale structural information.

## 1. INTRODUCTION

Fluorescence single-molecule localization microscopy (SMLM)^1–6^ can achieve spatial resolution beyond the diffraction limit of light by localizing the emitted light from many different fluorophores in acquired image frames. SMLM relies on sparse emitter concentrations where the emitter density in each frame must be controlled during the acquisition using imaging techniques such as (fluorescence) photoactivated localization microscopy ((f)PALM)^1,2^, (direct) stochastic optical reconstruction microscopy ((d)STORM)^3,7,4^, and DNA points accumulation in nanoscale topography (PAINT)^8^. Since most biological samples are inherently extended in 3D, the extension of SMLM from 2D to 3D is imperative for complete understanding of subcellular structures or biomolecular dynamics. Several techniques have been developed to obtain 3D coordinates of emitters, including multifocus^9–12^, interferometric^13^, and intensity-sensing methods^14^. Another approach to extract 3D information is using PSF engineering, which involves phase modulation of the light in the detection path of the microscope to encode both the lateral and axial position of the emitter into the shape of the PSF. The astigmatic PSF is a commonly used engineered PSF to obtain 3D super-resolution reconstructions over an axial range of ∼1 µm^15^. Multiple other engineered PSFs have been developed, including the corkscrew^16^, self-bending^17^, saddle-point^18^, tetrapod^19–21^, and nebulae PSFs^22,23^. The double-helix PSFs (DH-PSFs) have been extensively used for cellular super-resolution imaging^21,24–36^ and single-particle tracking (SPT)^37–41^ because of their relative ease of analysis and robustness for imaging at depths. The DH-PSFs are formed by a superposition of Gauss-Laguerre modes, and the resulting PSF consists of two lobes that revolve around their midpoint upon changes in the axial position of the emitter, appearing as a double-helix shape along the z-axis of the microscope^42,43^. The DH-PSF shape can be fit reasonably well to a double Gaussian function, where the midpoint between the lobes yields the lateral position of the emitter and the angle between the lobes yields the axial position. The DH phase patterns can be generated using either transmissive dielectric phase masks or a spatial light modulator.

Considerations when choosing an engineered PSF for 3D imaging include its achievable localization precision, effective axial range, and lateral footprint. PSFs with a short axial range typically offer good localization precision near the focal plane, but their performance usually deteriorates significantly outside of their designed axial range. SMLM imaging with short axial-range PSFs thus typically requires scanning over multiple, overlapping image slices to cover an entire cell volume^21,33,44,45^, which increases the complexity of both the experimental and analysis workflows, where the acquired image slices must be stitched together in post-processing to generate the complete 3D super-resolution reconstruction. Long axial-range PSFs, on the other hand, offer relatively uniform localization precisions throughout their extended axial range, holding great promise for imaging thick samples with a single slice. One drawback of long axial-range PSFs is that they often have large lateral footprints, which are more likely to cause overlapping emitters that are more challenging to localize with analysis algorithms relying on PSF fitting to model functions. To minimize the number of overlapping emitters, the image acquisitions therefore tend to be slower with long axial-range PSFs. However, this issue can be mitigated by using a matching pursuit-based method^46–48^ or a deep learning-based method such as DeepSTORM3D^23^ or DECODE^49^, which have been demonstrated to localize engineered PSFs in more densely emitting samples^33,23,49^.

Here, we demonstrate novel, long axial-range DH-PSFs for 3D super-resolution imaging of whole mammalian cells. We quantify the experimental localization precisions of DH-PSFs with 1-µm, 3-µm, 6-µm, and 12-µm axial ranges and the astigmatic PSF to benchmark their localization performance. Next, we demonstrate and quantitatively compare the performance of the 1-µm, 3-µm, and 6-µm axial range DH-PSFs for 3D SMLM of mammalian cells when analyzed using conventional PSF fitting. This approach allows for broad implementation by researchers who want to implement PSF engineering for stitching-free whole-cell imaging with a simple imaging and analysis pipeline. Finally, we show that using the deep learning-based analysis approach DeepSTORM3D for localization of the 6-µm axial range DH-PSF allows imaging with a higher emitter density and an up to 9 times higher image acquisition speed while achieving a resolution for scan-free whole-cell imaging below 30 nm laterally and 40 nm axially.

## 2. MATERIALS and METHODS

### 2.1. Experimental Setup

A schematic of the optical setup is shown in Figure 1a. A wide-field epi-fluorescence setup was built around an inverted microscope (IX83, Olympus) equipped with a 1.45 NA 100X oil immersion objective lens (UPLXAPO100X, Olympus). The excitation laser beams (560 nm, 2RU-VFL-P-1000-560-FCAPC, 1W; 642 nm: 2RU-VFL-P-1000-642-FCAPC, 1W, both MPB Communications) were circularly polarized (linear polarizers: LPVISC050-MP2, Thorlabs; and quarter-wave plates: 560 nm: Z-10-A-.250-B-556; 642 nm: Z-10-A-.250-B-647, both Tower Optical), spectrally filtered (560 nm: FF01-554/23-25; 642 nm: FF01-631/36-25, both Semrock), and collimated and expanded by lens telescopes (LA1951-A, *f* = 25.4 mm; LA1417-A, *f* = 150 mm, both Thorlabs) before being merged by a dichroic mirror (T5901pxr-UF2, Chroma) into a joint beam path. The beam was then further expanded using a lens telescope (AC508-075-A, *f* = 75 mm; AC508-300-A, *f* = 300 mm, both Thorlabs) and introduced to the back aperture of the objective lens through a Köhler lens (AC508-300-A, *f* = 300 mm, Thorlabs) to produce wide-field epi-illumination.

**Figure 1.**
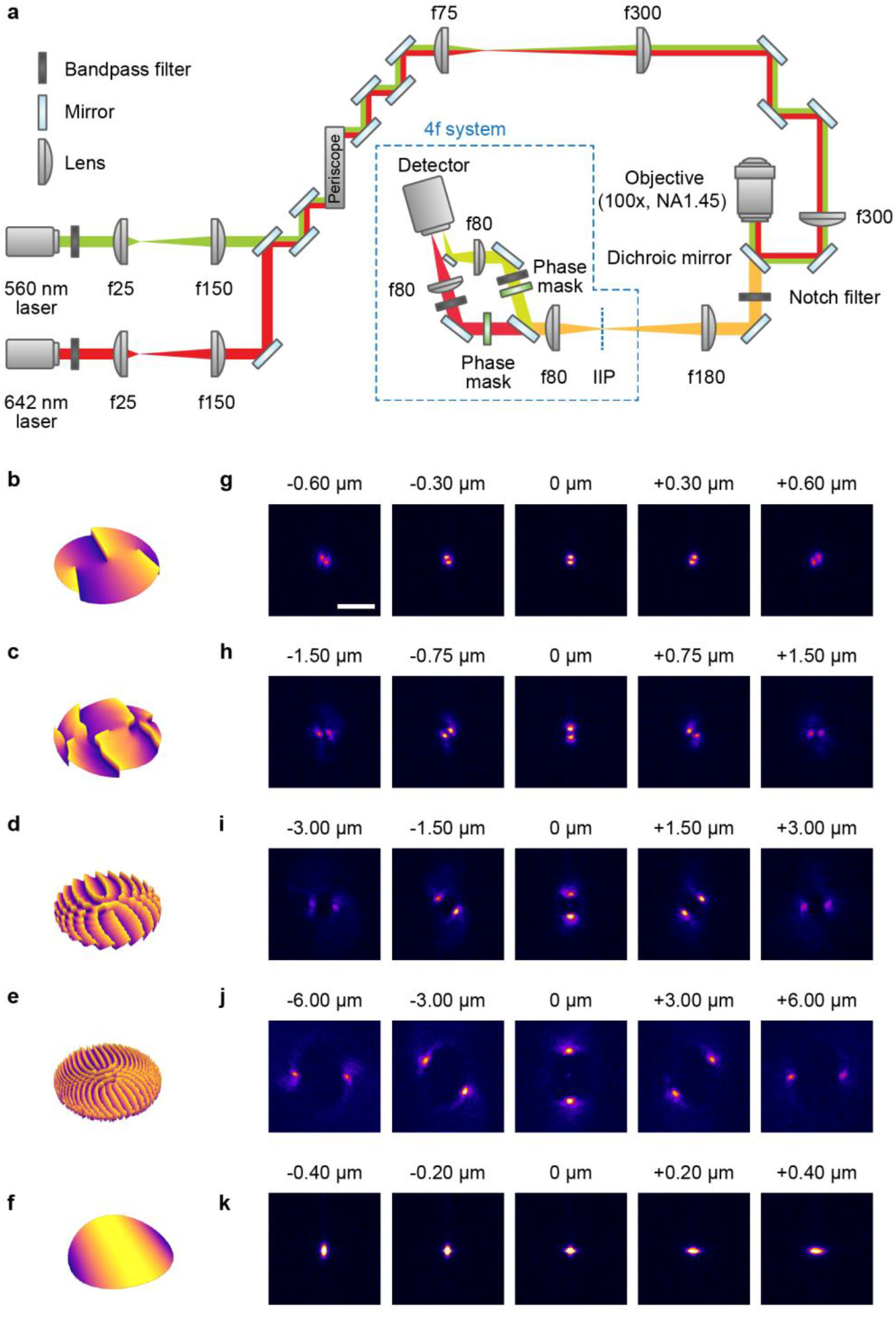
Design and characterization of the optical setup. (a) Schematic of the optical setup for 3D single-molecule localization microscopy with double-helix (DH) or astigmatic point spread functions (PSFs). The imaging system was built around a wide-field epi-fluorescence microscope equipped with a high-NA, oil-immersion objective lens. The detection path includes a two-color 4*f* system to access the Fourier plane of the microscope for PSF engineering. The schematic is not drawn to scale. (b)-(f) Schematic representations of the phase patterns of the transmissive dielectric DH phase masks with axial ranges of (b) 1 µm, (c) 3 µm, (d) 6 µm, and (e) 12 µm, and (f) of the astigmatic PSF. (g)-(k) Images of fluorescent beads acquired with DH-PSFs with axial ranges of (g) 1 µm, (h) 3 µm, (i) 6 µm, and (j) 12 µm, and (k) with the astigmatic PSF at the indicated axial positions. Scale bar 5 µm.

The samples were mounted on an xy translational stage (M26821LOJ, Physik Instrumente) and an xyz piezoelectric stage (OPH-PINANO-XYZ, Physik Instrumente).

The light emitted from the sample was collected by the objective lens, spectrally filtered (ZT405/488/561/640rpcV3 3 mm thick dichroic mirror in a Chroma BX3 cube; notch filters: ZET642NF, ZET561NF; all Chroma), and focused by the microscope tube lens to an image at the intermediate image plane (IIP), from which a two-channel 4*f* system was aligned in order to access the Fourier plane of the microscope for PSF engineering. The first lens of the 4*f* system (*f* = 80 mm, AC508-080-AB, Thorlabs) was positioned one focal length from the IIP. The light path was then split into two color-channels (termed “red” and “green”, respectively) using a dichroic mirror (T660lpxr-UF3, Chroma). Transmissive dielectric DH phase masks with 1-µm, 3-µm, 6-µm, or 12-µm axial range (red: DH2-670, DH1-670, DH6R-670, or DH12R-680, respectively; green: DH2-580, DH1-580, or DH6R-580, respectively, all Double Helix Optics Inc) were then positioned in the Fourier plane, one focal length from the first 4*f* lens, for PSF engineering (Figure 1b-e). The phase masks were mounted on magnetic mounts for easy placement and removal. To facilitate alignment, the magnetic mounts were placed on translational stages (red: PT3; green: PT1, both Thorlabs). The light was then further spectrally filtered (red: ET700/75; green: ET605/70m, both Chroma) and focused by second 4*f* lenses (*f* = 80 mm, AC508-080-AB, Thorlabs) positioned one focal length from the Fourier plane in each channel. The two light paths were then merged using a D-shaped mirror and imaged on different regions of an electron-multiplying CCD (EMCCD) camera (iXon Ultra 897, Andor). For the case of the astigmatic PSF, a weak cylindrical lens (LJ1516RM-A, *f* = 1000 mm, Thorlabs) was instead placed ∼10 mm after the second 4*f* lens, introducing astigmatism in the PSF shape.

### 2.2. Cell Culture

Human osteosarcoma cells (U-2 OS – HTB-96, ATCC) were plated on 8-well chambered glass coverslips (80827, Ibidi) and cultured at 37°C and 5% CO_2_ in Dulbecco’s Modified Eagle Medium (DMEM) (21063029, Gibco) supplemented with 10% (v/v) fetal bovine serum (FBS) (16000044, Gibco), and 100 mM sodium pyruvate (11360070, Gibco).

### 2.3. Sample Preparation

For PSF calibration and localization precision quantification, 0.1 µm fluorescent beads (Tetraspeck, T7279, Invitrogen) diluted 1:100 in nanopure water were diluted 1:10 in 10% polyvinyl alcohol (Mowiol 4-88, 17951, Polysciences Inc.) in nanopure water and spin coated (WS-650, Laurell Technologies) on glass coverslips (#1.5H, 22 × 22 mm, 170 ± 5 µm, 0102052, Marienfeld) which had been plasma cleaned with argon gas for 15 minutes using a plasma cleaner ((PDC-32G, Harrick Plasma).

For DNA-PAINT imaging of lamin B1, U-2 OS cells were fixed for 20 minutes at room temperature (RT) in 4% (w/v) formaldehyde solution made by diluting 16% formaldehyde solution (15710, Electron Microscopy Science) in 1X phosphate buffered saline (PBS) (SH3025602, Cytiva). The cells were then washed once with PBS and incubated in 10 mM ammonium chloride (213330, Sigma-Aldrich) in PBS for 10 minutes at RT. Next, the cells were permeabilized in 0.5% (v/v) Tween 20 (H5152, Promega) in PBS three times for 5 minutes at RT. The cells were then washed twice in ice-cold PBS and blocked for 30 minutes in PBS containing 2% donkey serum (ab7475, Abcam), 300 mM glycine (50046, Sigma-Aldrich), 0.05% (v/v) Tween 20, 0.1% (v/v) Triton X-100 (X100, Sigma-Aldrich), and 0.1% (w/v) BSA (A2058, Sigma-Aldrich). Next, the cells were labeled with rabbit anti-lamin B1 primary antibodies (ab16048, Abcam, 1:1000) diluted in incubation buffer (10 mM glycine, 0.05% (v/v) Tween 20, 0.1% Triton X-100, and 0.1% hydrogen peroxide (H325, Fisher Scientific) in PBS) for 1 hour at RT, then washed three times with 0.1% (v/v) Triton X-100 in PBS and once with washing buffer (Massive-AB-1-Plex, Massive Photonics) diluted 1:10 in nanopure water. The cells were then labeled with donkey anti-rabbit secondary antibodies conjugated with oligonucleotides (Massive-AB-1-Plex, Massive Photonics, diluted 1:100 in PBS with 0.1% (v/v) Tween 20) for 1 hour at RT. Then, the cells were washed twice with washing buffer before being incubated with 0.1 µm 580/605 nm fluorescent beads (F8801, Invitrogen) diluted 1:100,000 in nanopure water. After approximately 2 minutes of incubation, the cells were vigorously washed three times for 5 minutes with washing buffer and washed once in 500 mM sodium chloride (S9625-5006, Sigma-Aldrich) in PBS. The sample was then kept at 4°C until imaged. Finally, imager strands consisting of Cy3B conjugated with complementary oligonucleotides (Massive-AB-1-Plex, Massive Photonics) diluted in 500 mM sodium chloride in PBS were added to the sample at 0.2 nM.

For DNA-PAINT imaging of Tomm20, U-2 OS cells were fixed and quenched with ammonium chloride as described for the lamin B1 samples. The cells were permeabilized in 0.1% (w/v) saponin (SAE0073, Sigma Aldrich) for 10 minutes at RT and blocked with 10% (v/v) donkey serum (ab7475, Abcam), 0.1% (w/v) saponin, and 0.05 mg/mL salmon sperm ssDNA (ab229278, Abcam) in PBS for 1 hour at RT. The cells were then labeled with rabbit anti-Tomm20 primary antibodies (ab186735, Abcam) diluted 1:200 in 10% (v/v) donkey serum, 0.1% (w/v) saponin, and 0.05 mg/mL salmon sperm ssDNA in PBS for 1 hour at RT and washed three times with PBS for 5 minutes and once with washing buffer. The cells were then labeled with donkey anti-rabbit secondary antibodies which had been conjugated in house with azide-modified oligonucleotides with a sequence of TTATACATCTA (order No. 2023-67432, Biosynthesis) using dibenzocyclooctyne-sulfo-*N*-hydroxysuccinimidyl ester (762040, Sigma) as a cross-linker, using a previously described protocol^50^. The conjugated secondary antibodies were used for labeling at a dilution of 1:100 in antibody incubation buffer (Massive-AB-1-Plex, Massive Photonics) for 1 hour at RT. The cells were then washed three times with PBS for 7 minutes, three times with the washing buffer for 7 minutes, and once with PBS for 5 minutes. Samples were stored at 4°C until imaged. Before imaging, imager strands consisting of Cy3B conjugated with complementary oligonucleotides with a sequence of CTAGATGTAT (order No. 2023-67432, Biosynthesis) diluted in 500 mM sodium chloride in PBS were added to the sample at 0.25 nM.

### 2.4. Imaging Procedures and Settings

For all imaging, a set EM gain of 200 was used, which corresponded to a calibrated EM gain of 182 for the EMCCD camera. The EMCCD camera conversion gain was found experimentally to be 4.41 photoelectrons per A/D count. The camera was operated at a shift speed of 3.3 μm/s and normal vertical clock voltage amplitude. The read-out rate was set to 17 MHz at 16 bits using a preamplifier gain of 3. The calibrated pixel size of the camera was 159 nm/pixel vertically and 157 nm/pixel horizontally.

Calibration scans of each PSF were acquired using the piezoelectric xyz stage to translate the sample in the axial direction every five frames with step sizes of 250 nm for the 1-µm and 3-µm DH-PSF, 500 nm for the 6-µm DH-PSF, and 1000 nm for the 12-µm DH-PSF, and every frame with step size of 10 nm for the astigmatic PSF, with scan ranges of ±0.75 µm, ±1.5 µm, ±3 µm, ±6 µm, and ±0.75 µm for the scans with the 1-µm, 3-µm, 6-µm, and 12-µm DH-PSF and the astigmatic PSF, respectively.

To quantify and compare the experimental localization precisions of the different PSFs, a DH phase mask with an axial range of either 1 µm, 3 µm, 6 µm, or 12 µm (DH2-670, DH1-670, DH6R-670, or DH12R-680, respectively, Double Helix Optics Inc) or the cylindrical lens was placed in the red path in the optical setup. 500 frames of the beads were then acquired at each z position using step sizes and scan ranges of 0.1 µm and ±0.6 µm for the 1-µm DH-PSF, 0.25 µm and ±1.5 µm for 3-µm DH-PSF, 0.5 µm and ±2.5 µm for the 6-µm DH-PSF, 1 µm and ±6 µm for the 12-µm DH-PSF, and 0.1 µm and ±0.4 µm for the astigmatic PSF, with an exposure time of 50 ms using the 642 nm laser at intensities of ∼94 W/cm^2^, ∼27 W/cm^2^, and ∼8.1 W/cm^2^. These laser intensities were chosen to yield similar photon counts as expected from e.g., bright particles for SPT (∼15,000 photons per localization), intermediated photon counts comparable to e.g. DNA-PAINT and dSTORM imaging (∼6,000 photons per localization), and low photon counts comparable to e.g. imaging of fluorescent proteins (∼1,200 photons per localization). The number of frames per z position and the number of z-steps were selected to minimize the effects of photobleaching during the acquisitions. Beads were positioned in the center of the FOV for all measurements to avoid any field-dependent effects.

For sparse single-molecule imaging of lamin B1, a 1-µm, 3-µm, or 6-µm axial range DH phase mask (DH2-580, DH1-580, or DH6R-580, respectively, Double Helix Optics Inc) was placed in the Fourier plane of the green detection path. The sample was excited by the 560 nm laser at ∼210 W/cm^2^ with an exposure time of 250 ms. 50,000 frames were acquired for the 1-µm and 3-µm DH-PSF data, and 200,000 frames were acquired for the 6-µm DH-PSF data. Fiducial beads (F13082, Invitrogen) were imaged in the same channel to allow for drift correction in post-processing.

For dense single-molecule imaging of Tomm20, the 6-µm axial range DH phase mask (DH6R-580, Double Helix Optics Inc) was placed in the Fourier plane of the green detection path. The sample was excited by the 560 nm laser at ∼500 W/cm^2^ with an exposure time of 250 ms. 80,000 frames were acquired. Fiducial beads (F13082, Invitrogen) were imaged in the same channel to allow for drift correction in post-processing.

### 2.5. Data Analysis

The DH-PSF localization analyses were performed with the ImageJ plugin 3DTRAX® (Double Helix Optics Inc)^48^, or DeepSTORM3D^23^, a python-based open-source software for analyzing high-density datasets using a convolutional neural network (CNN).

For DH-PSF calibration in 3DTRAX, the bead images were cropped to include a single bead in a field of view (FOV) and then converted into hyperstacks in ImageJ before being loaded into 3DTRAX. The two lobes of the DH-PSF were fit with a double Gaussian function to determine the angle of the lobes at each known z position, yielding the calibration curves (Figure S1). For the astigmatic PSF calibration, the ImageJ plugin ThunderSTORM^51^ was used to calculate the calibration curve (Figure S2).

For bead localization precision measurements, the bead images acquired with the DH-PSFs were localized in 3DTRAX using the “sparse emitter” method and a detection threshold of 0.3-0.4. Bead data acquired using the astigmatic PSF were localized in ThunderSTORM using the “local maximum” detection method and the “elliptical Gaussian” localization method. After obtaining the localizations of the bead samples, drift correction was performed as described previously^20^ using a custom-written MATLAB code (Figure S3). In brief, the bead trajectories in each set of 500 frames were fitted to a third-degree polynomial function and subtracted from the original trajectory data. After the bead localization and drift correction process, experimental localization precisions were extracted by fitting the histograms of the localization distributions to Gaussian functions.

For CNN training using DeepSTORM3D, the phase pattern of the 6-µm axial range DH phase mask was retrieved from bead images using the VIPR software^52^. The recovered phase pattern was then used to simulate training data of the 6-µm axial range DH-PSF using an emitter density of 8 to 35 per 19.2 × 19.2 µm^2^ FOV, signal counts following a gamma distribution with a sharp parameter of 3 and a scale parameter of 3000, a minimal detectable signal counts of 2,500 counts per localization, and read noise in the range of 8 to 12, which matched the experimental single-molecule data from DNA-PAINT imaging of Tomm20.

For localizing single-molecule data, datasets from sparse emitter density DNA-PAINT imaging of lamin B1 using the 1-µm, 3-µm, and 6-µm axial range DH-PSFs were analyzed using 3DTRAX, and the mitochondria datasets acquired with the 6-µm axial range DH-PSF at high emitter densities were localized using both 3DTRAX and DeepSTORM3D. The fiducial bead was analyzed separately from the cell data using the same software as used for the single-molecule data. The analyzed fiducial bead localizations were then imported into a custom-written MATLAB script, where the trajectory of the fiducial bead was smoothed with a cubic spline and then subtracted from the single-molecule data to correct for drift. After drift correction, Vutara SRX (Bruker) was used to visualize the localizations from each dataset using a particle size of 20 nm. Lamin B1 data acquired with the 1-µm axial range DH-PSF was filtered to remove localizations with less than 2,000 photons per localization. Lamin B1 data acquired with the 3-µm and 6-µm axial range DH-PSFs were filtered to remove localizations with less than 1,000 photons per localization. The mitochondria data analyzed with 3DTRAX was filtered to remove localizations with less than 1,000 photons per localization. The mitochondria data analyzed with DeepSTORM3D was filtered for a global threshold less than 60. The mitochondria data analyzed with both approaches were filtered to remove localizations based on distance to nearest neighbors using a distance of 10 nm with 8 nearest neighbors. After filtering, Fourier ring correlation (FRC) analysis was performed using Vutara SRX to obtain the resulting resolution in xy, yz, and xz using a fixed threshold of 1/7 and a super-resolution pixel size of 8 nm.

## 3. RESULTS and DISCUSSION

### 3.1. Design and Characterization of the Optical Setup

To obtain calibration models for the DH-PSFs and the astigmatic PSF, images of fluorescent beads were acquired with the 1-µm, 3-µm, 6-µm, and 12-µm axial range DH-PSFs and the astigmatic PSF using transmissive dielectric DH phase masks or using the cylindrical lens (Figure 1). Calibration curves of the PSFs (Figures S1 and S2) were later used to localize the positions of emitters.

### 3.2. Quantification of the Localization Precision of Double-Helix Point Spread Functions of Varying Axial Range

The localization performances of the DH-PSFs with 1-µm, 3-µm, 6-µm, and 12-µm axial range and the astigmatic PSF were benchmarked by imaging of fluorescent beads. These measurements showed that the 3-µm axial range DH-PSF offers the best localization precision of the tested PSFs for high photon counts (∼15,000 photons per localization) (Figure 2a-c). The long (6-µm and 12-µm) axial-range DH-PSFs offer uniform localization performance across their respective axial ranges but at overall increased localization precision. A similar trend can be seen for intermediate photon counts (∼6,000 photons per localization), where the 3-µm axial range DH-PSF still exhibits the best localization precision (Figure 2d-f). For the case of low photon counts (∼1,200 photons per localization), the 1-µm axial range DH-PSF shows the best localization performance among the five PSFs (Figure 2g-i). The short axial-range DH-PSFs show superior performance compared to the conventional astigmatic PSF for all the used photon levels over comparable axial ranges. The localization precisions of the long axial-range DH-PSFs significantly deteriorate at these low photon counts, narrowing their effective detection range.

**Figure 2.**
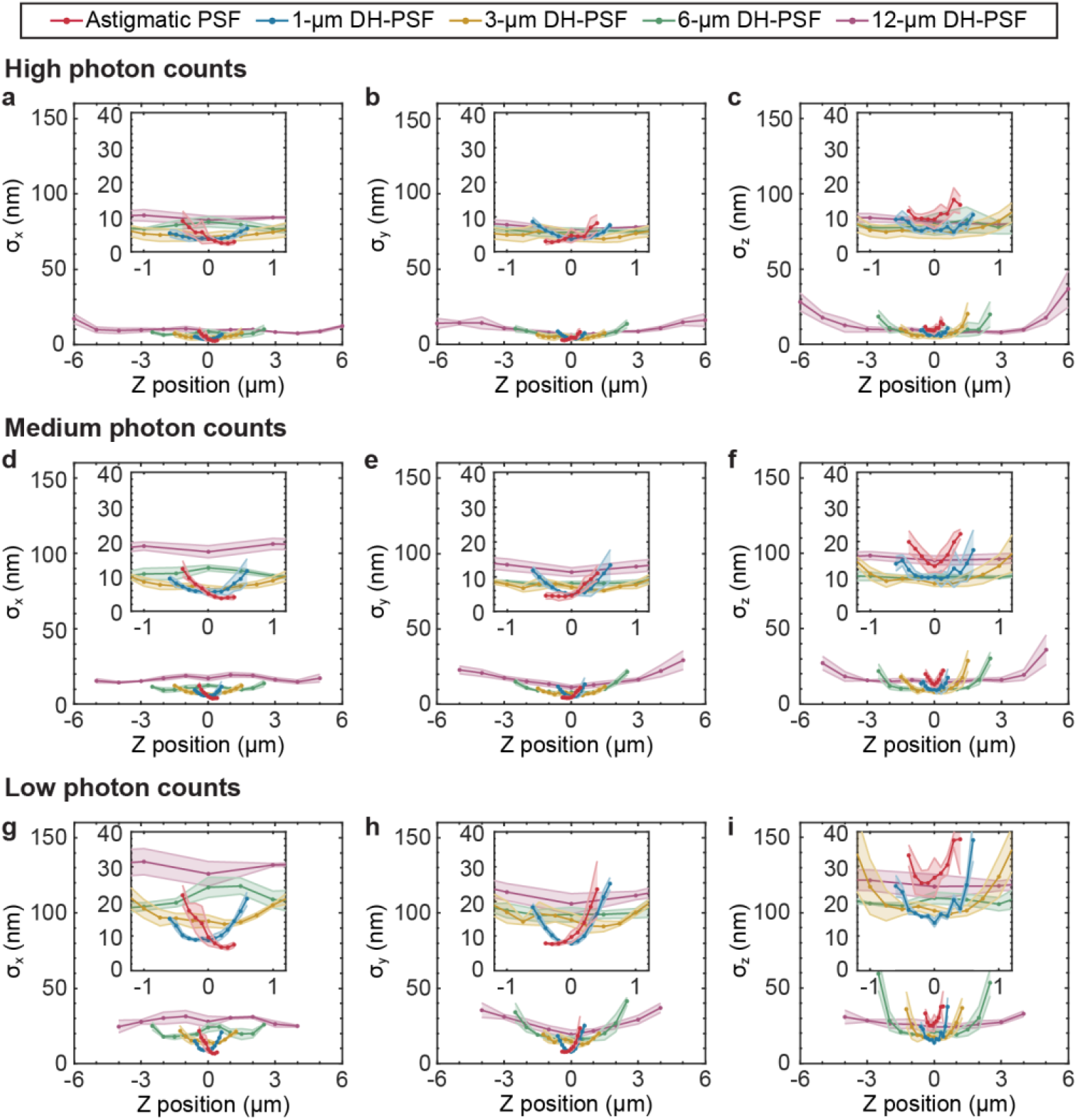
Quantification of the experimental localization precisions of double-helix (DH) point spread functions (PSFs) and the astigmatic PSF. (a)-(c) Localization precisions measured with a laser intensity of ∼94 W/cm^2^ in (a) x, (b) y, and (c) z. (d)-(f) Localization precisions measured with a laser intensity of ∼27 W/cm^2^ in (d) x, (e) y, and (f) z. (g)-(i) Localization precisions measured with a laser intensity of ∼8.1 W/cm^2^ in (g) x, (h) y, and (i) z. Points represent the mean values and the shaded areas represent the standard deviations from three independent measurements. Inserts in each figure show the localization precisions near the focal plane. Note that the axes of the inserts are on a different scale to highlight and compare the performance over a shorter axial range.

### 3.3. Comparison of the Performance of Double-Helix Point Spread Functions for Cellular 3D Single-molecule Super-resolution Imaging

To benchmark the performance of the DH-PSFs for 3D single-molecule super-resolution imaging of mammalian cells, we performed DNA-PAINT imaging of the nuclear lamina protein lamin B1 in U-2 OS cells using the 1-µm, 3-µm, and 6-µm axial range DH-PSFs. The resulting super-resolution reconstructions demonstrate the advantage of using the 6-µm axial range DH-PSF for capturing the entire nucleus without the need for stitching of slices (Figure 3a-c). FRC analysis was then used to estimate the achievable resolution using the three DH-PSFs (Figure 3d-f). The 1-µm DH-PSF offers the best FRC resolutions (28.6/35.3/32.5 nm in the xy/xz/yz plane) among the three DH-PSFs, resulting in ∼293,550 localizations from 50,000 frames (Figure 3d). However, it only allows capture of less than 1-µm axial range in a single slice (Figure 3a). Using the 3-µm DH-PSF allowed reconstruction of the lamina structure with ∼522,520 localizations from 50,000 frames over a ∼2.1 µm axial range (Figure 3b) with FRC values of 43.8/45.8/48.9 nm in xy/xz/yz (Figure 3e), showing cellular features including a nuclear lamina channel and a nuclear invagination. The entire nuclear lamina structure was reconstructed over a ∼3.5-µm axial range with ∼439,180 localizations from 200,000 frames with a single slice of imaging data acquired with the 6-µm DH-PSF (Figure 3c) with FRC values of 60.4/65.9/64.3 nm in xy/xz/yz (Figure 3f), demonstrating scan-free 3D super-resolution imaging of the whole mammalian cell nucleus.

**Figure 3.**
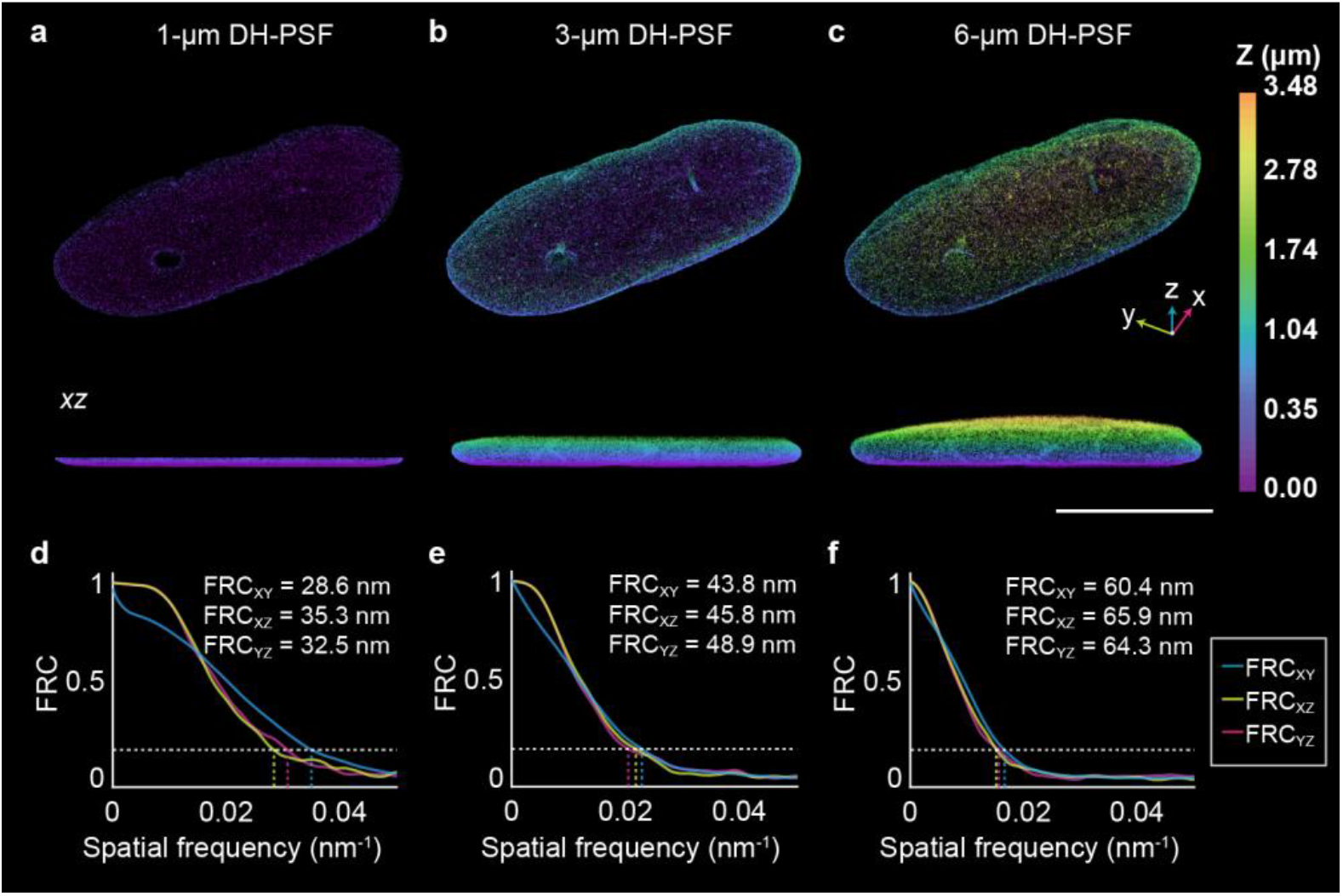
Comparison of the performance of double-helix (DH) point spread functions (PSFs) for 3D single-molecule super-resolution imaging of the nuclear lamina. (a)-(c) Super-resolution reconstructions of lamin B1 acquired using DH-PSFs with axial ranges of (a) 1 µm, (b) 3 µm, and (c) 6 µm. (d)-(f) Fourier ring correlation (FRC) curves in the xy, xz, and yz planes of the data shown in (a), (b), and (c), respectively. Scale bar 10 µm.

### 3.4. Speed and Resolution Improvement of Scan-free Whole-cell 3D Single-Molecule Super-resolution Imaging by Deep-learning based Analysis

To demonstrate and quantify the improvement in acquisition speed by deep-learning based analysis, 3D single-molecule imaging of mitochondria was performed using the 6-µm axial range DH-PSF at high emitter densities and the data was analyzed using DeepSTORM3D and 3DTRAX (Figure 4 and Figure S4). While analysis with 3DTRAX, which relies on conventional PSF fitting to a double Gaussian function, only localized an average of ∼6 emitters per frame (a total of ∼480,000 localizations in 80,000 frames) when using a detection threshold of 0.3, DeepSTORM3D detected an average of ∼29 localizations per frame (a total of ∼2,320,000 localizations in 80,000 frames), resulting in a ∼5-fold improvement in acquisition speed when using DeepSTORM3D for the whole cell sample. The resulting FRC resolutions of the whole-cell mitochondria data reconstructed with DeepSTORM3D (Figure 4a) were 26.2/34.0/36.0 nm in xy/xz/yz compared to 44.3/46.7/46.5 nm in xy/xz/yz when using 3DTRAX (Figure 4b) for analysis of this dense data. The result demonstrates the overall resolution improvement for the same number of frames when using DeepSTORM3D while allowing for ∼5 times faster data acquisition speed. The difference in acquisition speed is even more dramatic when considering denser regions of the sample, where DeepSTORM3D could improve the acquisition speed by up to 9 times while still improving the FRC values compared to 3DTRAX (Figure 4c-d).

**Figure 4.**
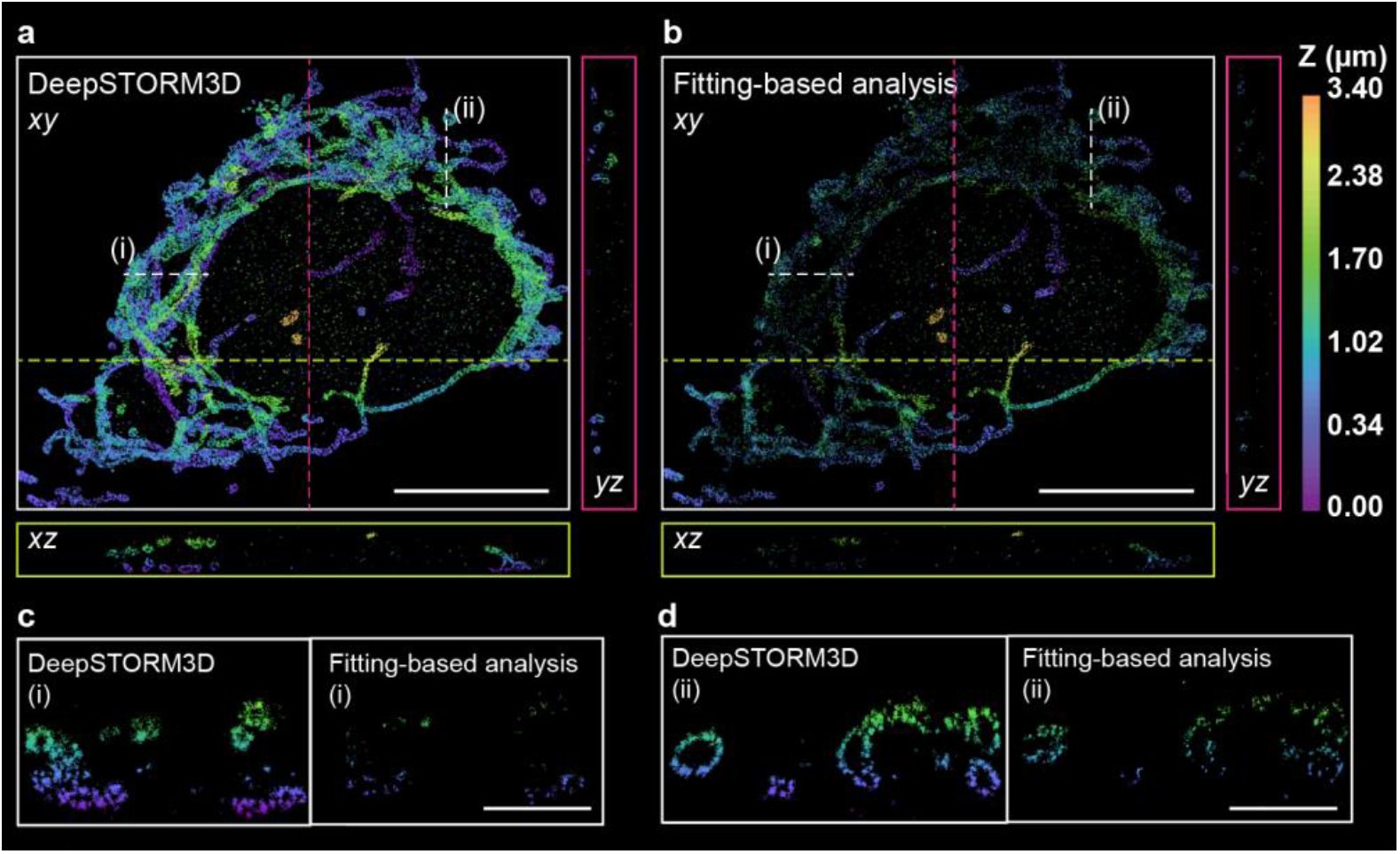
Speed and resolution improvement by DeepSTORM3D for 3D single-molecule super-resolution whole-cell imaging of mitochondria. (a, b) Whole-cell 3D super-resolution reconstruction of Tomm20 acquired with the 6-µm axial range DH-PSF at dense emitter concentration and analyzed using (a) the deep-learning based approach DeepSTORM3D and (b) fitting-based analysis using 3DTRAX. The xz and yz views show 300-nm thick xz and yz cross-sections along the dashed magenta lines and yellow lines shown in the xy view, respectively. Scale bars 10 µm. (c) 300 nm thick xz cross-sections along the dashed line (i) in (a) and (b). The FRC resolution of the cross-sections were found to be 41.6 nm and 46.6 nm while yielding a total number of ∼10,300 and 1,200 localizations for the DeepSTORM3D data and the 3DTRAX data, respectively. Scale bar 2 µm. (d) 300 nm thick yz cross-sections along the dashed line (ii) in (a) and (b). FRC resolutions of the cross-sections were found to be 32.7 nm and 34.6 nm while yielding a total number of 9,900 and 2,500 localizations for the DeepSTORM3D data and the 3DTRAX data, respectively. Scale bar 2 µm.

## 4. CONCLUSIONS

Our experimental localization precision measurements demonstrate that the short axial-range DH-PSFs show superior performance compared to the conventional astigmatic PSF for all the used photon levels over comparable axial ranges. The 3-µm axial range DH-PSF offers similar localization performance as the 1-µm axial range DH-PSF for high and medium photon levels, but over a larger axial range. The 1-µm axial range DH-PSF becomes superior when localizing emitters at low photon levels. The DH-PSFs with long axial ranges (6 µm and 12 µm) offer uniform localization performance across their axial ranges, providing the benefit of simpler analysis than stitching of multiple slices acquired with a shorter-range PSF for imaging of thick samples. This was further demonstrated by 3D single-molecule imaging of the nuclear lamina in mammalian cells, where the 6-µm axial range DH-PSF allowed scan-free 3D super-resolution imaging across the whole sample volume with a single slice, simplifying the experimental and analysis workflows. Furthermore, integrating DeepSTORM3D into the localization analysis for high emitter density datasets improved the image acquisition speed by up to a factor of 9 while resulting in scan-free whole cell imaging with a resolution below 40 nm in all dimensions.

These long axial-range PSFs are compatible with other commonly used SMLM imaging methods, such as dSTORM and PALM, and can be applied for multi-target imaging by using Exchange-PAINT. Furthermore, the localization performance of long axial-range DH-PSFs in SMLM can be improved by using selective illumination, such as light-sheet illumination^21,33^, or by using self-quenching DNA imager strands^53^ to minimize the fluorescence background. Spherical aberration correction using e.g. adaptive optics^54^ will further improve the localization performance, especially when imaging at depth.

In summary, we here demonstrate that SMLM with long axial-range DH-PSFs allows scan-free, whole-cell, 3D single-molecule super-resolution imaging, offering a simpler imaging and analysis workflow for whole-cell samples and with drastically improved image acquisition speed and resolution when utilizing deep-learning based analysis.

## ASSOCIATED CONTENT

### Supporting Information

See supplemental document for supplemental figures associated with this work.

### Data Availability

Data presented in this paper can be obtained from the authors upon reasonable request.

### Code Availability

Astigmatic PSF data was analyzed using the open-source ImageJ plugin ThunderSTORM^51^ (https://github.com/zitmen/thunderstorm). Calibration and localization analysis of DH-PSF data were performed using 3DTRAX^48^. Localization analysis of dense 6-µm DH-PSF data was performed using DeepSTORM3D^23^ (https://github.com/EliasNehme/DeepSTORM3D). The custom-written MATLAB codes for drift correction and localization precision estimation are available upon reasonable request.

## AUTHOR INFORMATION

### Author Contribution

The manuscript was written through contributions of all authors. All authors have given approval to the final version of the manuscript.

## ACKNOWLEDGEMENTS

The authors thank Warren Colomb, Nahima Saliba, Gabriella Gagliano, and Sofia Vargas-Hernandez for helpful discussions and Margareth Freire for help labeling the Tomm20 samples. The authors thank Elias Nehme and Dafei Xiao for help with neural network training and localization analysis with DeepSTORM3D.

## REFERENCES

(1) Betzig, E.; Patterson, G. H.; Sougrat, R.; Lindwasser, O. W.; Olenych, S.; Bonifacino, J. S.; Davidson, M. W.; Lippincott-Schwartz, J.; Hess, H. F. Imaging Intracellular Fluorescent Proteins at Nanometer Resolution. Science 2006, 313 (5793), 1642–1645. 10.1126/science.1127344.

(2) Hess, S. T.; Girirajan, T. P. K.; Mason, M. D. Ultra-High Resolution Imaging by Fluorescence Photoactivation Localization Microscopy. Biophys. J. 2006, 91 (11), 4258– 4272. 10.1529/biophysj.106.091116.

(3) Rust, M. J.; Bates, M.; Zhuang, X. Sub-Diffraction-Limit Imaging by Stochastic Optical Reconstruction Microscopy (STORM). Nat Methods 2006, 3 (10), 793–795. 10.1038/nmeth929.

(4) Heilemann, M.; van de Linde, S.; Schüttpelz, M.; Kasper, R.; Seefeldt, B.; Mukherjee, A.; Tinnefeld, P.; Sauer, M. Subdiffraction-Resolution Fluorescence Imaging with Conventional Fluorescent Probes. Angew. Chem. Int. Ed. 2008, 47 (33), 6172–6176. 10.1002/anie.200802376.

(5) Gustavsson, A.-K.; Petrov, P. N.; Moerner, W. E. Light Sheet Approaches for Improved Precision in 3D Localization-Based Super-Resolution Imaging in Mammalian Cells [Invited]. Opt. Express 2018, 26 (10), 13122. 10.1364/OE.26.013122.

(6) Gagliano, G.; Nelson, T.; Saliba, N.; Vargas-Hernández, S.; Gustavsson, A.-K. Light Sheet Illumination for 3D Single-Molecule Super-Resolution Imaging of Neuronal Synapses. Front. Synaptic Neurosci. 2021, 13, 761530. 10.3389/fnsyn.2021.761530.

(7) Bates, M.; Huang, B.; Dempsey, G. T.; Zhuang, X. Multicolor Super-Resolution Imaging with Photo-Switchable Fluorescent Probes. Science 2007, 317 (5845), 1749–1753. 10.1126/science.1146598.

(8) Jungmann, R.; Avendaño, M. S.; Woehrstein, J. B.; Dai, M.; Shih, W. M.; Yin, P. Multiplexed 3D Cellular Super-Resolution Imaging with DNA-PAINT and Exchange-PAINT. Nat Methods 2014, 11 (3), 313–318. 10.1038/nmeth.2835.

(9) Ram, S.; Prabhat, P.; Chao, J.; Sally Ward, E.; Ober, R. J. High Accuracy 3D Quantum Dot Tracking with Multifocal Plane Microscopy for the Study of Fast Intracellular Dynamics in Live Cells. Biophys. J. 2008, 95 (12), 6025–6043. 10.1529/biophysj.108.140392.

(10) Hajj, B.; Wisniewski, J.; El Beheiry, M.; Chen, J.; Revyakin, A.; Wu, C.; Dahan, M. Whole-Cell, Multicolor Superresolution Imaging Using Volumetric Multifocus Microscopy. Proc. Natl. Acad. Sci. USA 2014, 111 (49), 17480–17485. 10.1073/pnas.1412396111.

(11) Toprak, E.; Balci, H.; Blehm, B. H.; Selvin, P. R. Three-Dimensional Particle Tracking via Bifocal Imaging. Nano Lett. 2007, 7 (7), 2043–2045. 10.1021/nl0709120.

(12) Abrahamsson, S.; Chen, J.; Hajj, B.; Stallinga, S.; Katsov, A. Y.; Wisniewski, J.; Mizuguchi, G.; Soule, P.; Mueller, F.; Darzacq, C. D.; Darzacq, X.; Wu, C.; Bargmann, C. I.; Agard, D. A.; Dahan, M.; Gustafsson, M. G. L. Fast Multicolor 3D Imaging Using Aberration-Corrected Multifocus Microscopy. Nat. Methods 2013, 10 (1), 60–63. 10.1038/nmeth.2277.

(13) Huang, F.; Sirinakis, G.; Allgeyer, E. S.; Schroeder, L. K.; Duim, W. C.; Kromann, E. B.; Phan, T.; Rivera-Molina, F. E.; Myers, J. R.; Irnov, I.; Lessard, M.; Zhang, Y.; Handel, M. A.; Jacobs-Wagner, C.; Lusk, C. P.; Rothman, J. E.; Toomre, D.; Booth, M. J.; Bewersdorf, J. Ultra-High Resolution 3D Imaging of Whole Cells. Cell 2016, 166 (4), 1028–1040. 10.1016/j.cell.2016.06.016.

(14) Dickson, R. M.; Norris, D. J.; Tzeng, Y.-L.; Moerner, W. E. Three-Dimensional Imaging of Single Molecules Solvated in Pores of Poly(Acrylamide) Gels. Science 1996, 274 (5289), 966–968. 10.1126/science.274.5289.966.

(15) Huang, B.; Wang, W.; Bates, M.; Zhuang, X. Three-Dimensional Super-Resolution Imaging by Stochastic Optical Reconstruction Microscopy. Science 2008, 319 (5864), 810– 813. 10.1126/science.1153529.

(16) Lew, M. D.; Lee, S. F.; Badieirostami, M.; Moerner, W. E. Corkscrew Point Spread Function for Far-Field Three-Dimensional Nanoscale Localization of Pointlike Objects. Opt. Lett. 2011, 36 (2), 202. 10.1364/OL.36.000202.

(17) Jia, S.; Vaughan, J. C.; Zhuang, X. Isotropic Three-Dimensional Super-Resolution Imaging with a Self-Bending Point Spread Function. Nat. Photonics 2014, 8 (4), 302–306. 10.1038/nphoton.2014.13.

(18) Shechtman, Y.; Sahl, S. J.; Backer, A. S.; Moerner, W. E. Optimal Point Spread Function Design for 3D Imaging. Phys. Rev. Lett. 2014, 113 (13), 133902. 10.1103/PhysRevLett.113.133902.

(19) Shechtman, Y.; Weiss, L. E.; Backer, A. S.; Sahl, S. J.; Moerner, W. E. Precise Three-Dimensional Scan-Free Multiple-Particle Tracking over Large Axial Ranges with Tetrapod Point Spread Functions. Nano Lett. 2015, 15 (6), 4194–4199. 10.1021/acs.nanolett.5b01396.

(20) Shechtman, Y.; Gustavsson, A.-K.; Petrov, P. N.; Dultz, E.; Lee, M. Y.; Weis, K.; Moerner, W. E. Observation of Live Chromatin Dynamics in Cells via 3D Localization Microscopy Using Tetrapod Point Spread Functions. Biomed. Opt. Express 2017, 8 (12), 5735. 10.1364/BOE.8.005735.

(21) Gustavsson, A.-K.; Petrov, P. N.; Lee, M. Y.; Shechtman, Y.; Moerner, W. E. 3D Single-Molecule Super-Resolution Microscopy with a Tilted Light Sheet. Nat Commun 2018, 9 (1), 123. 10.1038/s41467-017-02563-4.

(22) Nehme, E.; Ferdman, B.; Weiss, L. E.; Naor, T.; Freedman, D.; Michaeli, T.; Shechtman, Y. Learning Optimal Wavefront Shaping for Multi-Channel Imaging. IEEE Transactions on Pattern Analysis and Machine Intelligence 2021, 43 (7), 2179–2192. 10.1109/TPAMI.2021.3076873.

(23) Nehme, E.; Freedman, D.; Gordon, R.; Ferdman, B.; Weiss, L. E.; Alalouf, O.; Naor, T.; Orange, R.; Michaeli, T.; Shechtman, Y. DeepSTORM3D: Dense 3D Localization Microscopy and PSF Design by Deep Learning. Nat Methods 2020, 17 (7), 734–740. 10.1038/s41592-020-0853-5.

(24) Pavani, S. R. P.; Thompson, M. A.; Biteen, J. S.; Lord, S. J.; Liu, N.; Twieg, R. J.; Piestun, R.; Moerner, W. E. Three-Dimensional, Single-Molecule Fluorescence Imaging beyond the Diffraction Limit by Using a Double-Helix Point Spread Function. Proc. Natl. Acad. Sci. USA 2009, 106 (9), 2995–2999. 10.1073/pnas.0900245106.

(25) Grover, G.; DeLuca, K.; Quirin, S.; DeLuca, J.; Piestun, R. Super-Resolution Photon-Efficient Imaging by Nanometric Double-Helix Point Spread Function Localization of Emitters (SPINDLE). Opt. Express, OE 2012, 20 (24), 26681–26695. 10.1364/OE.20.026681.

(26) Gahlmann, A.; Ptacin, J. L.; Grover, G.; Quirin, S.; von Diezmann, A. R. S.; Lee, M. K.; Backlund, M. P.; Shapiro, L.; Piestun, R.; Moerner, W. E. Quantitative Multicolor Subdiffraction Imaging of Bacterial Protein Ultrastructures in Three Dimensions. Nano Lett. 2013, 13 (3), 987–993. 10.1021/nl304071h.

(27) Carr, A. R.; Ponjavic, A.; Basu, S.; McColl, J.; Santos, A. M.; Davis, S.; Laue, E. D.; Klenerman, D.; Lee, S. F. Three-Dimensional Super-Resolution in Eukaryotic Cells Using the Double-Helix Point Spread Function. Biophysical Journal 2017, 112 (7), 1444–1454. 10.1016/j.bpj.2017.02.023.

(28) Möckl, L.; Pedram, K.; Roy, A. R.; Krishnan, V.; Gustavsson, A.-K.; Dorigo, O.; Bertozzi, C. R.; Moerner, W. E. Quantitative Super-Resolution Microscopy of the Mammalian Glycocalyx. Dev. Cell 2019, 50 (1), 57–72. 10.1016/j.devcel.2019.04.035.

(29) Bayas, C. A.; Diezmann, A. von; Gustavsson, A.-K.; Moerner, W. E. Easy-DHPSF 2.0: Open-Source Software for Three-Dimensional Localization and Two-Color Registration of Single Molecules with Nanoscale Accuracy. Prot. Exch. 2019. 10.21203/rs.2.9151/v2.

(30) Bennett, H. W.; Gustavsson, A.-K.; Bayas, C. A.; Petrov, P. N.; Mooney, N.; Moerner, W. E.; Jackson, P. K. Novel Fibrillar Structure in the Inversin Compartment of Primary Cilia Revealed by 3D Single-Molecule Superresolution Microscopy. Mol. Biol. Cell 2020, 31 (7), 619–639. 10.1091/mbc.E19-09-0499.

(31) Sanders, E. W.; Carr, A. R.; Bruggeman, E.; Körbel, M.; Benaissa, S. I.; Donat, R. F.; Santos, A. M.; McColl, J.; O’Holleran, K.; Klenerman, D.; Davis, S. J.; Lee, S. F.; Ponjavic, A. resPAINT: Accelerating Volumetric Super-Resolution Localisation Microscopy by Active Control of Probe Emission**. Angewandte Chemie International Edition 2022, 61 (42), e202206919. 10.1002/anie.202206919.

(32) Kanie, T.; Love, J. F.; Fisher, S. D.; Gustavsson, A.-K.; Jackson, P. K. A Hierarchical Pathway for Assembly of the Distal Appendages That Organize Primary Cilia. bioRxiv 2023, 2023.01.06.522944. 10.1101/2023.01.06.522944.

(33) Gagliano, G.; Saliba, N.; Gustavsson, A.-K. Whole-Cell Multi-Target Single-Molecule Super-Resolution Imaging in 3D with Microfluidics and a Single-Objective Tilted Light Sheet. bioRxiv 2023, 2023.09.27.559876. 10.1101/2023.09.27.559876.

(34) Weiss, L. E.; Love, J. F.; Yoon, J.; Comerci, C. J.; Milenkovic, L.; Kanie, T.; Jackson, P. K.; Stearns, T.; Gustavsson, A.-K. Chapter 4 - Single-Molecule Imaging in the Primary Cilium. In Methods in Cell Biology; Bravo-San Pedro, J.M., Galluzzi, L., Eds.; Academic Press, 2023; Vol. 176, pp 59–83. 10.1016/bs.mcb.2023.01.003.

(35) Chowdhury, P.; Wang, X.; Love, J. F.; Vargas-Hernandez, S.; Nakatani, Y.; Grimm, S. L.; Mezquita, D.; Mason, F. M.; Martinez, E. D.; Coarfa, C.; Walker, C. L.; Gustavsson, A.-K.; Dere, R. Lysine Demethylase 4A Is a Centrosome Associated Protein Required for Centrosome Integrity and Genomic Stability. bioRxiv 2024, 2024.02.20.581246. 10.1101/2024.02.20.581246.

(36) Nelson, T.; Vargas-Hernández, S.; Freire, M.; Cheng, S.; Gustavsson, A.-K. Multimodal Illumination Platform for 3D Single-Molecule Super-Resolution Imaging throughout Mammalian Cells. Biomed. Opt. Express 2024, 15 (5), 3050–3063. 10.1364/BOE.521362.

(37) Thompson, M. A.; Casolari, J. M.; Badieirostami, M.; Brown, P. O.; Moerner, W. E. Three-Dimensional Tracking of Single mRNA Particles in Saccharomyces Cerevisiae Using a Double-Helix Point Spread Function. Proceedings of the National Academy of Sciences 2010, 107 (42), 17864–17871. 10.1073/pnas.1012868107.

(38) Yu, B.; Yu, J.; Li, W.; Cao, B.; Li, H.; Chen, D.; Niu, H. Nanoscale Three-Dimensional Single Particle Tracking by Light-Sheet-Based Double-Helix Point Spread Function Microscopy. Appl. Opt. 2016, 55 (3), 449. 10.1364/AO.55.000449.

(39) Gustavsson, A.-K.; Ghosh, R. P.; Petrov, P. N.; Liphardt, J. T.; Moerner, W. E. Fast and Parallel Nanoscale Three-Dimensional Tracking of Heterogeneous Mammalian Chromatin Dynamics. Mol. Biol. Cell 2022, 33 (6), 1–11. 10.1091/mbc.E21-10-0514.

(40) Wang, F.; Lai, J.; Liu, H.; Zhao, M.; Zhang, Y.; Xu, J.; Yu, Y.; Wang, C. Double Helix Point Spread Function with Variable Spacing for Precise 3D Particle Localization. Opt. Express 2023, 31 (7), 11680–11694. 10.1364/OE.482390.

(41) Basu, S.; Shukron, O.; Hall, D.; Parutto, P.; Ponjavic, A.; Shah, D.; Boucher, W.; Lando, D.; Zhang, W.; Reynolds, N.; Sober, L. H.; Jartseva, A.; Ragheb, R.; Ma, X.; Cramard, J.; Floyd, R.; Balmer, J.; Drury, T. A.; Carr, A. R.; Needham, L.-M.; Aubert, A.; Communie, G.; Gor, K.; Steindel, M.; Morey, L.; Blanco, E.; Bartke, T.; Di Croce, L.; Berger, I.; Schaffitzel, C.; Lee, S. F.; Stevens, T. J.; Klenerman, D.; Hendrich, B. D.; Holcman, D.; Laue, E. D. Live-Cell Three-Dimensional Single-Molecule Tracking Reveals Modulation of Enhancer Dynamics by NuRD. Nat Struct Mol Biol 2023, 30 (11), 1628–1639. 10.1038/s41594-023-01095-4.

(42) Piestun, R.; Schechner, Y. Y.; Shamir, J. Propagation-Invariant Wave Fields with Finite Energy. J. Opt. Soc. Am. A, JOSAA 2000, 17 (2), 294–303. 10.1364/JOSAA.17.000294.

(43) Pavani, S. R. P.; Piestun, R. High-Efficiency Rotating Point Spread Functions. Opt. Express 2008, 16 (5), 3484. 10.1364/OE.16.003484.

(44) Huang, B.; Jones, S. A.; Brandenburg, B.; Zhuang, X. Whole-Cell 3D STORM Reveals Interactions between Cellular Structures with Nanometer-Scale Resolution. Nat. Methods 2008, 5 (12), 1047–1052. 10.1038/nmeth.1274.

(45) Rozario, A. M.; Morey, A.; Elliott, C.; Russ, B.; Whelan, D. R.; Turner, S. J.; Bell, T. D. M. 3D Single Molecule Super-Resolution Microscopy of Whole Nuclear Lamina. Frontiers in Chemistry 2022, 10.

(46) Barsic, A.; Grover, G.; Piestun, R. Three-Dimensional Super-Resolution and Localization of Dense Clusters of Single Molecules. Sci Rep 2014, 4 (1), 5388. 10.1038/srep05388.

(47) Babcock, H. P.; Zhuang, X. Analyzing Single Molecule Localization Microscopy Data Using Cubic Splines. Sci Rep 2017, 7 (1), 552. 10.1038/s41598-017-00622-w.

(48) Gaumer, S.; Colomb, W.; Loiacono, A.; Kimerling, L.; Agrawal, A. Computational Recovery of Engineered Point Spread Functions in Single Molecule Localization Microscopy Using the Double Helix 3DTRAX Software. Microscopy and Microanalysis 2021, 27 (S1), 868–871. 10.1017/S143192762100338X.

(49) Speiser, A.; Müller, L.-R.; Hoess, P.; Matti, U.; Obara, C. J.; Legant, W. R.; Kreshuk, A.; Macke, J. H.; Ries, J.; Turaga, S. C. Deep Learning Enables Fast and Dense Single-Molecule Localization with High Accuracy. Nat Methods 2021, 18 (9), 1082–1090. 10.1038/s41592-021-01236-x.

(50) Schnitzbauer, J.; Strauss, M. T.; Schlichthaerle, T.; Schueder, F.; Jungmann, R. Super-Resolution Microscopy with DNA-PAINT. Nat. Protoc. 2017, 12 (6), 1198–1228. 10.1038/nprot.2017.024.

(51) Ovesný, M.; Křížek, P.; Borkovec, J.; Švindrych, Z.; Hagen, G. M. ThunderSTORM: A Comprehensive ImageJ Plug-in for PALM and STORM Data Analysis and Super-Resolution Imaging. Bioinformatics 2014, 30 (16), 2389–2390. 10.1093/bioinformatics/btu202.

(52) Ferdman, B.; Nehme, E.; Weiss, L. E.; Orange, R.; Alalouf, O.; Shechtman, Y. VIPR: Vectorial Implementation of Phase Retrieval for Fast and Accurate Microscopic Pixel-Wise Pupil Estimation. Opt. Express 2020, 28 (7), 10179. 10.1364/OE.388248.

(53) Kessler, L. F.; Balakrishnan, A.; Deußner-Helfmann, N. S.; Li, Y.; Mantel, M.; Glogger, M.; Barth, H.-D.; Dietz, M. S.; Heilemann, M. Self-Quenched Fluorophore Dimers for DNA-PAINT and STED Microscopy. Angew. Chem. Int. Ed. 2023, 62 (39), e202307538. 10.1002/anie.202307538.

(54) Zhang, P.; Ma, D.; Cheng, X.; Tsai, A. P.; Tang, Y.; Gao, H.-C.; Fang, L.; Bi, C.; Landreth, G. E.; Chubykin, A. A.; Huang, F. Deep Learning-Driven Adaptive Optics for Single-Molecule Localization Microscopy. Nature Methods 2023, 20 (11), 1748–1758. 10.1038/s41592-023-02029-0.

