## Supporting Information for "Long axial-range double-helix point spread functions for 3D volumetric super-resolution imaging"

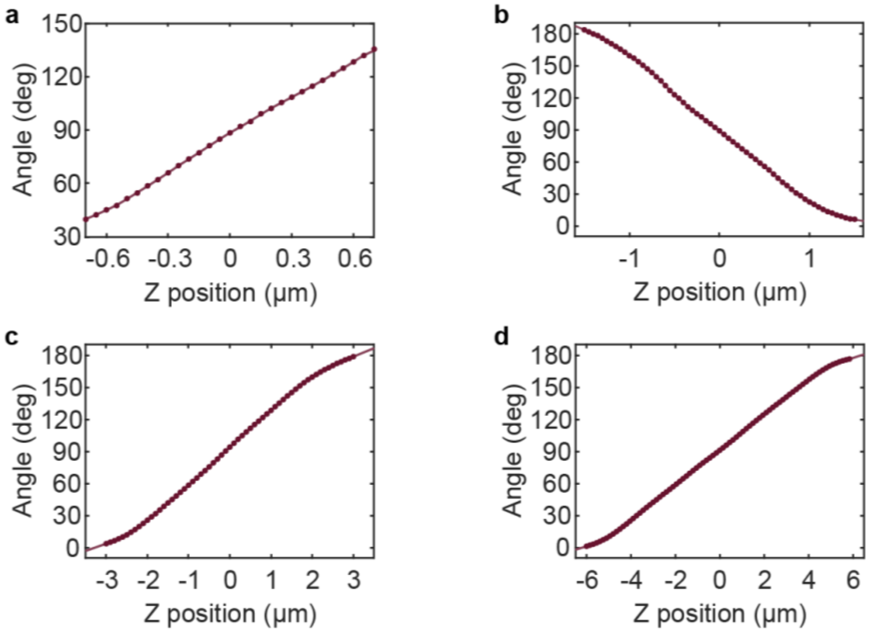


Figure S1. Representative calibration curves of double-helix point spread functions (DH-PSFs) with (a) 1-µm, (b) 3-µm, (c) 6-µm, and (d) 12-µm axial range.

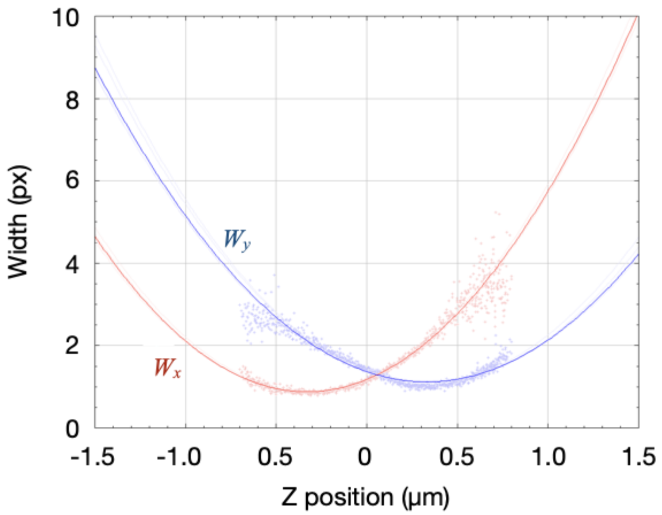


Figure S2. Representative calibration curve of the astigmatic PSF.

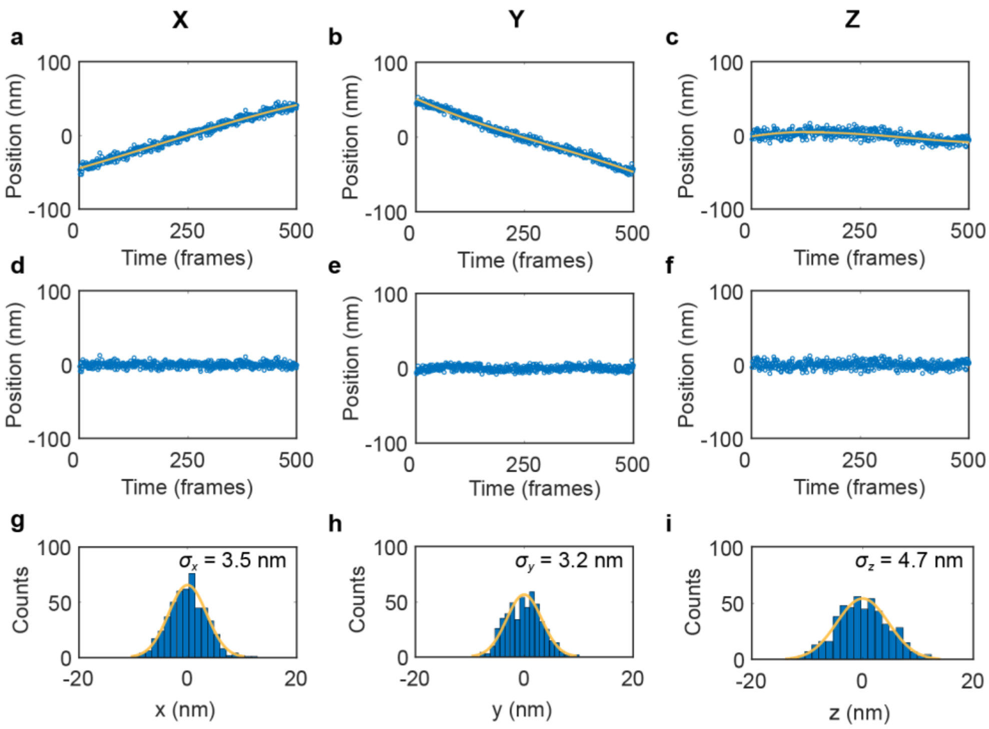


Figure S3. Experimental localization precision estimation from imaging of fluorescent beads. (a)-(c) Localized bead positions (blue rings) and third-degree polynomial fits (yellow lines) in (a) x, (b) y, and (c) z. (d)-(e) Localized bead positions after drift correction by subtracting the fitted values from the data in (d) x, (e) y, and (f) z. (g)-(i) Histograms showing localization distributions in (g) x, (h) y, and (i) z (blue bars) of the data shown in (d)-(f). Experimental localization precisions in each dimension were estimated from the standard deviations of Gaussian fits of the localized distribution (yellow line) after drift correction.

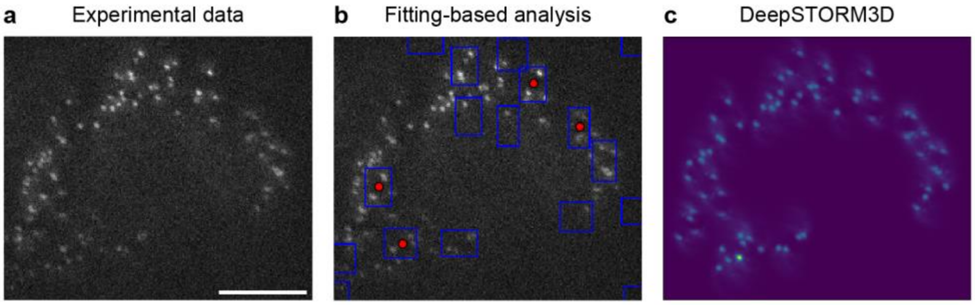


Figure S4. (a) Representative raw experimental image frame of the mitochondria single-molecule data acquired with the 6-µm axial range DH-PSF reconstructed in Fig. 4. (b) Localization results using fitting-based analysis (3DTRAX), demonstrating that four emitters were localized (red dots) in this frame. (c) Localization results using DeepSTORM3D for analysis, demonstrating that 34 emitters were detected in this frame. Scale bar 10 µm.
